# Does gender influence cardiovascular remodeling in C57BL/6J mice fed a high-fat, high-sucrose, high-salt diet?

**DOI:** 10.1101/367524

**Authors:** Debora Cristina Pereira-Silva, Rayane Paula Machado-Silva, Camila Castro-Pinheiro, Caroline Fernandes-Santos

## Abstract

Animal models are widely used to study the physiopathology of human diseases. However, the influence of gender on modern society diet style-induced cardiovascular disease was not exploited so far. Thus, this study investigated cardiovascular remodeling in C57BL/6J mice fed a diet rich in saturated fat, sucrose, and salt, evaluating gender effect on this process. Male and female C57BL/6J mice were fed AIN93M diet or a modified AIN93M rich in fat, sucrose, and salt (HFSS) for 12 weeks. Body mass, water and food intake and cardiovascular remodeling were assessed. The HFSS diet did not lead to body mass gain or glucose metabolism disturbance assessed by serum glucose, insulin, and oral glucose tolerance test. However, female mice on a HFSS diet had increased visceral and subcutaneous adiposity. Only male mice displayed heart hypertrophy. The left ventricle was not hypertrophied in male and female mice, but its lumen was dilated. Intramyocardial arteries and the thoracic aorta had intima-media thickening in male mice, but in the female, it was only noticed in the thoracic aorta. Finally, intramyocardial artery dilation was present in both genders, but not in the aorta. Changes in LV dimensions and the arterial remodeling were influenced by both gender and the HFSS diet. In conclusion, male and female C57BL/6J mice suffered cardiovascular remodeling after 12 weeks of high-fat, high-sucrose, high-salt feeding, although they did not develop obesity or diabetes. Sexual dimorphism occurred in response to diet for body adiposity, heart hypertrophy, and intramyocardial artery remodeling.

## Introduction

Animals models are widely used to study human diseases and are important tools to understand the physiopathology of several conditions such as cardiovascular disease (CVD) (Leong *et al*., 2015). Over the last decades, they have helped the development of drugs and new treatment strategies, since they function as a living system and respond to different approaches. However, much that is currently known was built based on male rodents while the female has been overlooked. For instance, sex-related differences in pharmacokinetics and pharmacodynamics exist, and it forces the study of gender differences on drug efficacy (Baggio *et al*., 2013).

Premenopausal women display a lower incidence of CVD compared to postmenopausal women and young adult men. It shows that estrogen ― or the lack of androgens ― might play a protective role in CVD development in premenopausal women (Deschepper e Llamas, 2007; Chakrabarti *et al*., 2014). Male and female present several differences regarding coronary artery size, electrophysiological heart properties, gene expression, contractile properties and how this organ responds to injury (Deschepper e Llamas, 2007). Gender impact on the occurrence, prognosis, and response to treatment of various CVD has recently been a focus of interest (Azevedo *et al*., 2016), which reinforces the need to understand how gender impact CVD development in animal models.

Diet is a risk factor for CVD, and it is attributed mainly to the metabolic impact of salt overload, and to sugar and fat quantity and quality (Aneja *et al*., 2004). Salt overload correlates with higher blood pressure levels, and it changes hemodynamic parameters (Van Huysse *et al*., 2011). Saturated fatty acids consumption is strongly correlated with increased CVD incidence, compensatory hyperinsulinemia, hyperglycemia, and hyperleptinemia (Martha et al., 2011). Finally, simple carbohydrates contribute to the high prevalence of obesity and type 2 diabetes, which also contributes to CV outcomes. Among carbohydrates, fructose is one of the main components of the modern occidental diet, especially sucrose, and its consumption is largely associated with insulin resistance and arterial hypertension (Te Morenga *et al*., 2014; Sanders, 2015).

In animal models, some CVD phenotypes may be induced by diet. Diets rich in sugar or fat, alone or in combination, lead to diabetes and obesity (Fernandes-Santos, Carneiro, *et al*., 2009; Costa *et al*., 2012; Sanders, 2015), whereas salt alone leads to hypertension and cardiovascular remodeling (Meneton *et al*., 2005; Aaron e Sanders, 2013). However, the combination of fat, sugar, and salt overload is less exploited in literature, and the sex response to these nutrients combined. Pigs fed a diet rich in fat, sucrose, and salt have increased blood pressure (Myrie *et al*., 2012). Similar reports in human studies indicate that salt prevents the development of obesity in men (Polonia e Martins, 2009; Ekinci *et al*., 2011).

Overall, it is extremely important to investigate sex-specific aspects of cardiovascular remodeling in rodent models of diet-induced CV remodeling, since it has not been deeply exploited so far for sugar, fat and salt overload. It represents the diet style of the modern society that has been contributing to the obesity epidemy. Thus, this study investigated cardiovascular remodeling in C57BL/6J mice fed a diet rich in saturated fat, sucrose, and salt, evaluating gender effect on this process.

## Material and methods

### Ethical aspects

The handling and experimental protocols were approved by the local Ethics Committee to Care and Use of Laboratory Animals (CEUA#647/15). The study was performed following the Animal Research Reporting In vivo Experiments ARRIVE guidelines and the Guideline for the Care and Use of Laboratory Animals (US NIH Publication N° 85–23. Revised 1996) (Kilkenny et al., 2010).

### Animals and diet

Male and female C57BL/6J mice at two months of age were obtained from colonies maintained at the Universidade Federal Fluminense and kept under standard conditions (collective polypropylene cages, 12h light/dark cycles, 21±2ºC, humidity 60±10% and air exhaustion cycle 15min/h). At three months old, mice were randomly allocated into four groups according to diet (n=12–15/group). Control group received AIN93M diet (Reeves *et al*., 1993), and the experimental groups received a high-fat high-sucrose high-salt diet (HFSS) modified from the AIN93M diet (Pragsolucoes, Jau, Sao Paulo, Brazil) for 12 weeks. The AIN93M diet consisted of 3.84 Kcal/g and 0.25% NaCl (w/w), and the HFSS diet consisted of 4.39 Kcal/g and 8% NaCl (w/w). The main difference is the addition of saturated fat (lard, 36.9% energy/kg), simple carbohydrates (sucrose, 27.3% energy/kg) and sodium chloride to the HFSS diet (table 1). For 12 weeks, food and water were offered ad libitum, and its ingestion monitored daily and weekly, respectively. Body mass was assessed weekly, and body fat evaluated at euthanasia by harvesting visceral and subcutaneous fat depots (genital and inguinal fat pads, respectively) (Shimadzu, AUW220D, Kyoto, Japan).

**Table 1.** Experimental diet

| **Ingredients** | **AIN93M** | **HFSS** |
| --- | --- | --- |
| **g/kg** |  |  |
| Casein | 140.0 | 160.0 |
| L-cystine | 1.8 | 1.8 |
| Corn starch | 620.7 | 140.7 |
| Sucrose | 100.0 | 300.0 |
| Fibers (cellulose) | 50.0 | 50.0 |
| Soy oil | 40,0 | 40.0 |
| Lard | - | 180.0 |
| Vitamin mix * | 10.0 | 10.0 |
| Mineral mix * | 35.0 | 35.0 |
| NaCl ** | - | 80.0 |
| Choline | 2.5 | 2.5 |
| Antioxidant | 0.008 | 0.06 |
| **Energy** |  |  |
| Kcal/g | 3.84 | 4.39 |
| Total protein, % | 14.8 | 14.7 |
| Total lipid, % | 9.4 | 45.1 |
| Total carbohydrate, % | 75.8 | 40.2 |

\* For vitamin and mineral mix composition, see Reeves et al., 1993.
\*\* AIN93M has 0,25% NaCl and the HFSS diet has 8% NaCl.

**Table 2.** Cardiovascular remodeling

| Parameter | Male |  | Female |  | Two-Way ANOVA |  |  |
| --- | --- | --- | --- | --- | --- | --- | --- |
|  | Control | HFSS diet | Control | HFSS diet | Gender | Diet | Interaction |
| *Biometry* |  |  |  |  |  |  |  |
| Heart/tibia, mg/cm | 7.72 ± 1.32 | 9.19 ± 0.66 * | 6.09 ± 0.48 # | 6.78 ± 0.52 # | < 0.0001 | 0.005 | NS |
| LV/tibia, mg/cm | 5.29 ± 0.94 | 5.92 ± 0.60 | 4.32 ± 0.34 | 4.45 ± 0.31 # | 0.0002 | NS | NS |
| LV/heart, mg/mg | 0.69 ± 0.021 | 0.66 ± 0.030 | 0.67 ± 0.054 | 0.66 ± 0.037 | NS | NS | NS |
| *Left ventricle* |  |  |  |  |  |  |  |
| Free wall, mm | 1.39 ± 0.11 | 1.25 ± 0.12 | 1.02 ± 0.14 # | 1.16 ± 0.16 | 0.0013 | NS | 0.03 |
| Interventricular septum, mm | 1.45 ± 0.09 | 1.58 ± 0.08 | 1.16 ± 0.33 | 1.30 ± 0.10 | 0.0019 | NS | NS |
| Lumen diameter, mm | 1.35 ± 0.07 | 1.86 ± 0.13 * | 0.95 ± 0.14 # | 1.27 ± 0.05 * # | < 0.0001 | < 0.0001 | 0.04 |
| Free wall: Lumen | 1.04 ± 0.12 | 0.67 ± 0.07 * | 1.07 ± 0.07 | 0.91 ± 0.13 # | 0.0084 | < 0.0001 | 0.03 |
| IV Septum: Lumen | 1.09 ± 0.12 | 0.85 ± 0.09 | 1.20 ± 0.18 | 1.02 ± 0.08 | 0.015 | 0.0011 | NS |
| *Intramyocardial arteries* |  |  |  |  |  |  |  |
| IMT, µm | 7.63 ± 1.54 | 14.68 ± 4.10 * | 8.32 ± 0.93 | 10.91 ± 1.82 | NS | 0.0003 | NS |
| Lumen diameter, µm | 44.1 ± 5.9 | 80.7 ± 30.1 * | 32.9 ± 3.7 | 70.0 ± 9.8 * | NS | < 0.0001 | NS |
| IMT: Lumen | 0.178 ± 0.052 | 0.197 ± 0.072 * | 0.256 ± 0.039 | 0.156 ± 0.015 | NS | NS | NS |
| *Thoracic aorta* |  |  |  |  |  |  |  |
| IMT, µm | 43.4 ± 0.41 | 55.2 ± 1.34 * | 43.6 ± 3.7 | 52.6 ± 0.59 * | NS | < 0.0001 | NS |
| Lumen diameter, µm | 522.2 ± 71.0 | 561.9 ± 45.3 | 450.8 ± 64.9 | 469.3 ± 18.6 # | 0.0012 | NS | NS |
| IMT: Lumen | 0.085 ± 0.013 | 0.099 ± 0.008 * | 0.089 ± 0.10 | 0.112 ± 0.004 * | 0.02 | < 0.0001 | NS |

Data are mean ± S.D., p<0.05, \* control (same gender), # male (similar diet). NS, non-significant. Abbreviations: HFSS, high-fat, high-sucrose, high-salt diet; LV, left ventricle; IV, interventricular; IMT, intima-media thickness.

### Glucose metabolism

On the day of euthanasia, mice were deeply anesthetized with ketamine 100.0mg/kg (Francotar®, Virbac, Brazil) and xylazine 10.0mg/kg (Virbaxyl 2%®, Virbac, Brazil) intraperitoneal, and the heart was exposed for blood collection from the right atrium. Serum was obtained by centrifugation (120g for 10min) and used to measure glucose and insulin concentrations (rat/mouse insulin ELISA #E6013-K, Millipore). Oral glucose tolerance test (OGTT) was performed at week 11. After a 6-hour fast, 50% glucose in sterile saline (0.9%NaCl) with a dose of 1g/kg was administered by orogastric gavage. Blood samples were milked from the tail tip by a small incision. Plasma glucose concentration was measured (glucometer Accu-Chek Performa Nano, Roche Diagnostic, Germany) before glucose gavage and later at 15, 30, 60 and 90 minutes after glucose administration. The area under the curve (AUC) was calculated using the trapezoid rule to assess glucose intolerance.

### Left ventricle remodeling

The heart was removed and weighed, and the left ventricle (LV) was carefully dissected and isolated for weighing, to evaluate cardiac and LV hypertrophy (Shimadzu, AUW220D, Kyoto, Japan). Left tibia length was measured from malleolus to medial condyle to correct cardiac and LV mass, to minimize the effect of animal size on these parameters (Yin *et al*., 1982). LV samples were immersed in 4% buffered formalin pH 7.2 for 48h, then cut transversally at their half height based on base-apex axis. The apex fragment undergone routine histological processing, embedded in Paraplast plus (P3683, Sigma), cut at 5μm thick, and stained with hematoxylin and eosin.

Morphometry focused on the geometric aspects of the LV chamber and its intramyocardial arteries (fig. 1a-d). Six animals per group and three non-consecutive sections per animal were used. Images of LV chamber were acquired at 4x magnification and arteries at 40x (Evos XL, ThermoScientific), and measurements were performed using the Image-Pro® Plus software v.4.5 (Media Cybernetics, Silver Spring, USA). It was assessed the lumen diameter (LD) of LV chamber, and the LV free wall (FW) and interventricular septum (IVS) thickness at the level of papillary muscles (Slawson *et al*., 1998). Intramyocardial artery LD and intima-media thickness (IMT) were also evaluated.

**Figure 1.**
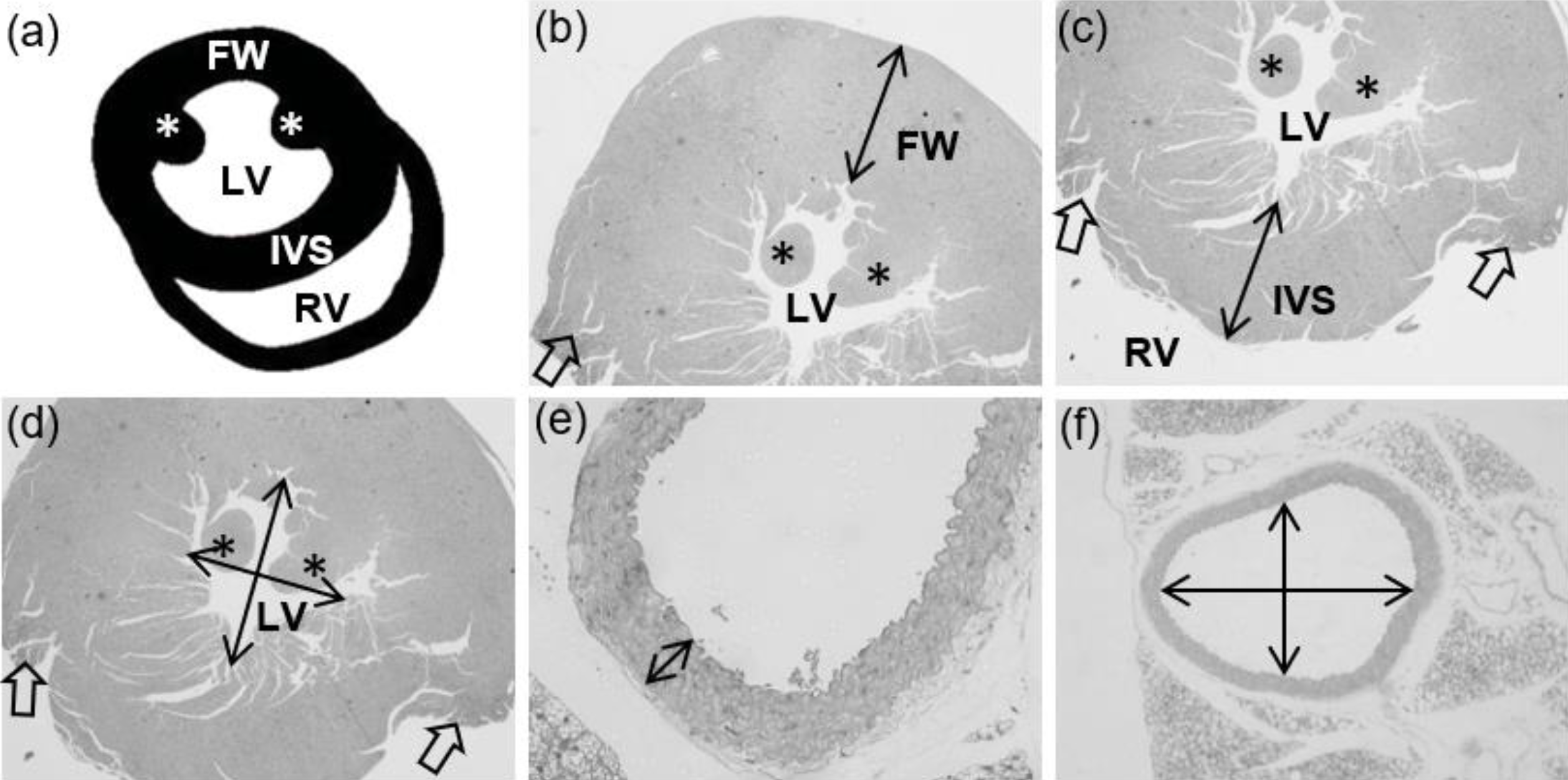
Cardiovascular morphometry. **a**, Scheme of the heart sectioned in the transverse plane, where the left (LV) and right ventricles (RV) lumen are in evidence, and the free wall (FW) and the interventricular septum (IVS) wall. The papillary muscles are indicated by an asterisk (*). Heart orientation matches photomicrographs in **b-d**, and it is not in anatomical orientation. **b-d**, Photomicrographs illustrating how the FW (**b**), IVS (**c**) and LV lumen diameter (**d**) were measured. Open arrows indicate the site where RV wall was dissected in euthanasia. **e-f**, Photomicrography is illustrating the measure of intima-media thickness (**e**) and lumen diameter (**f**) of the thoracic aorta in the transverse plane. The same was performed for intramyocardial arteries.

### Aorta remodeling

On the day of euthanasia, the thoracic aorta was harvested and followed the same routine histological procedures as previously mentioned for the LV. For morphometry, six animals were used per group, and three histological sections of the artery were produced per mice. Images were acquired at 4x magnification to assess LD and at 40x to assess IMT as previously described and shown in fig. 1e-f (Fernandes-Santos, De Souza Mendonça*, et al*., 2009).

### Statistics

Data are expressed as mean ± SD and analyzed for normality and homoscedasticity of variances. Comparison among groups was made by ANOVA two-way followed by a post-hoc test of Tukey. A *P*-value <0.05 was considered statistically significant (GraphPad® Prism software v. 6.0, La Jolla, CA, USA).

## Results

### Diet rich in fat, sucrose, and salt do not induce obesity or hyperglycemia in C57BL/6J mice

As expected, mice fed with the HFSS diet had increased water intake of about 156% for both male and female mice when compared to their respective control AIN93M diet groups (P<0.0001, fig. 2a). Although food intake was similar among groups, energy consumption was 20% and 14% higher in HFSS male and female mice, respectively, compared to control groups (P<0.01, fig. 2b-c). Water and energy intake were influenced by both gender and diet (two-way ANOVA).

**Figure 2.**
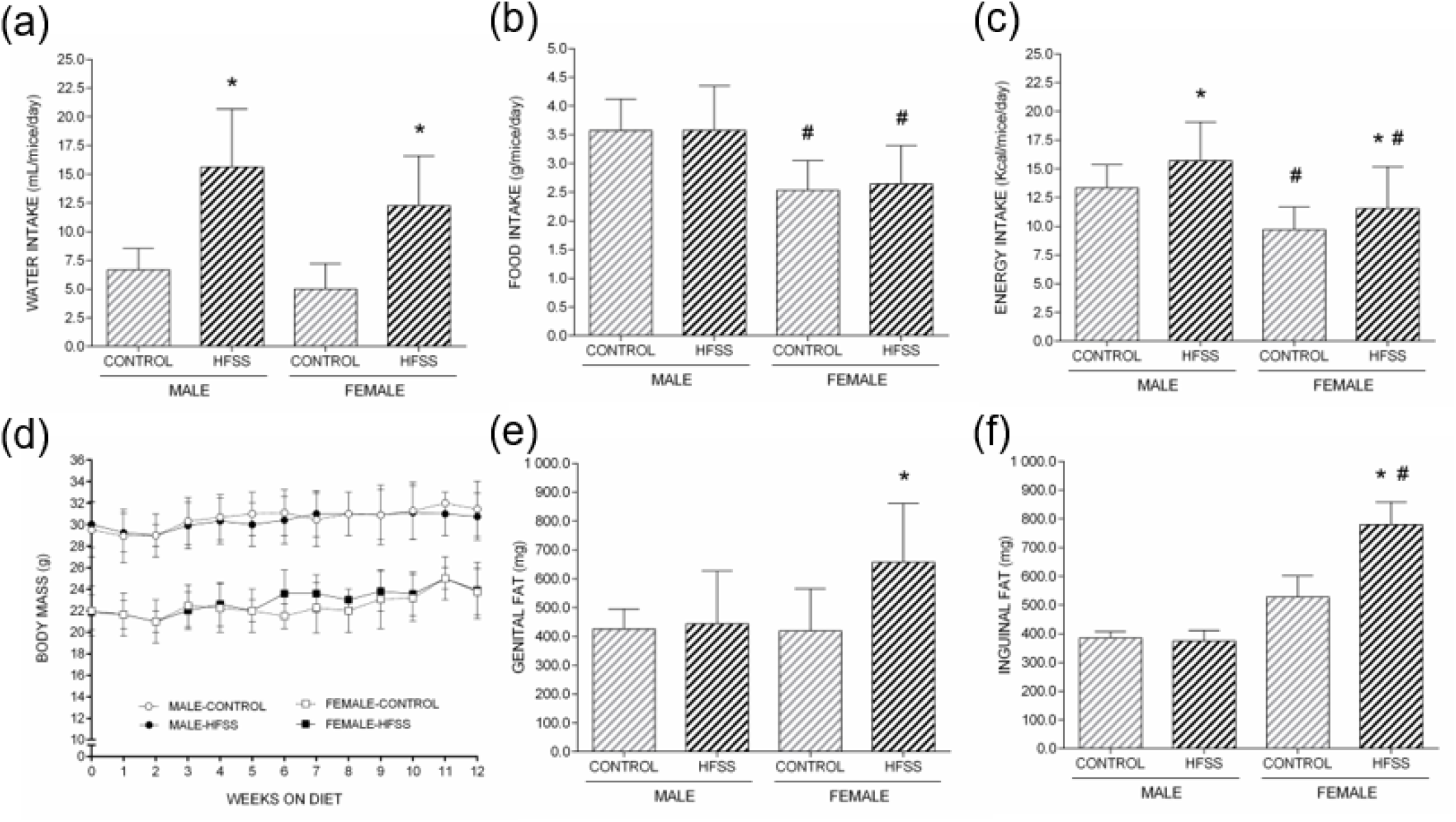
Water (**a**), food (**b**) and energy (**c**) consumption per mice daily. Body mass response to 12 weeks on high-fat, high-sucrose, high-salt (HFSS) diet in shown in (**d**), and body adiposity, represented by visceral (genital) and subcutaneous (inguinal) fat is shown in **e-f**. Data are mean ± S.D., p<0.05, * control (same gender), # male (similar diet).

An intriguing result was that HFSS mice did not gain weight along the twelve weeks of HFSS diet ingestion (fig. 2d). Nevertheless, female mice on HFSS diet presented increased body adiposity, characterized by 66% increase in visceral fat (genital, P<0.05) and 48% in subcutaneous fat (inguinal, P<0.05) depots (fig. 2e-f). The same was not true for male mice, that showed no increase of fat depots by the HFSS diet. The two-way ANOVA showed a gender effect on inguinal fat (P<0.001), but not on genital fat (P=0.099), which in turn displayed a minor influence by the diet (P<0.04).

Glucose metabolism was not disturbed in male mice by the HFSS diet since no significant changes were noticed for blood glucose, insulin or glucose tolerance (fig. 3). A similar result was found for female mice, but an intriguing finding was an improvement of glucose tolerance after 12 weeks on the HFSS diet (fig. 3c-d).

**Figure 3.**
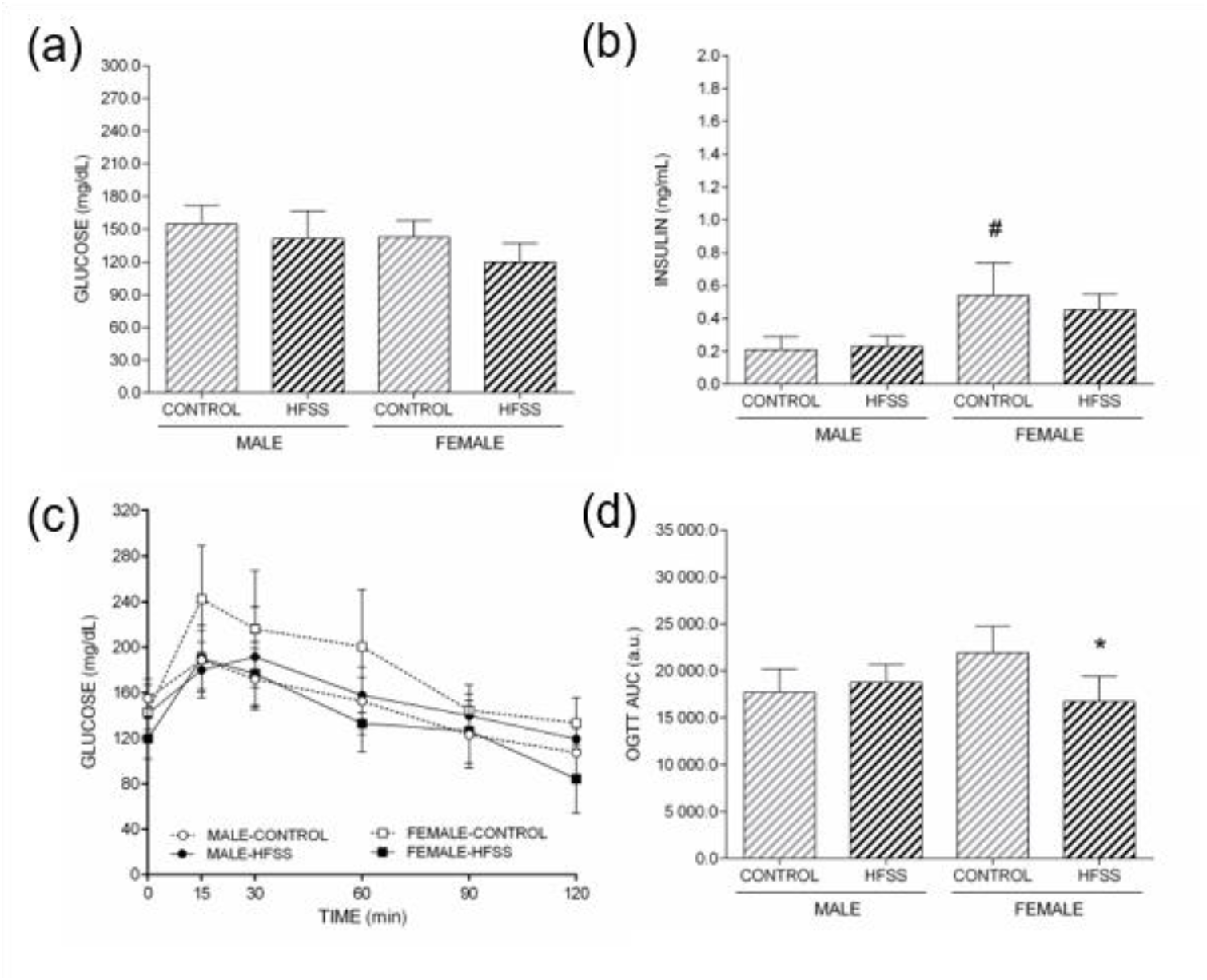
Serum glucose (**a**), and insulin (**b**). Oral glucose tolerance test (OGTT) curve is shown in (**c**) and its area under the curve (AUC) in (**d**). a.u., arbitrary units. Data are mean ± S.D., p<0.05, * control (same gender), # male (similar diet).

### Gender and diet differently modulates cardiovascular remodeling

Twelve weeks on HFSS diet lead to heart hypertrophy only in male mice (+19% heart/tibia ratio, P<0.05), and both gender and diet influenced this parameter (table 3). No significant change in LV mass was noticed when it was assessed the LV/tibia and LV/heart ratio (table 3). Morphometry of LV wall corroborates with this result since the free wall and the interventricular septum thickness was not increased in male and female mice on the HFSS diet (table 3), displaying a solely gender-dependent difference in size as expected (two-way ANOVA, table 3). On the other hand, the hyperenergetic and hypersodic diet lead to LV chamber dilation, since lumen diameter increased 38% (P<0.0001) in male and 34% (P<0.001) in female mice, a result of both diet and gender interaction (two-way ANOVA, table 3 and fig. 4a-d). Due to LV dilation, the free wall/lumen ratio in male mice fed with the HFSS diet presented a reduction of 36% (P<0.001, table 3). In summary, although some LV parameters did not present significant difference between control and HFSS groups, the twoway analysis of variance showed that LV wall dimension is dependent on gender, and lumen diameter is dependent on both gender and diet intake (table 3).

**Figure 4.**
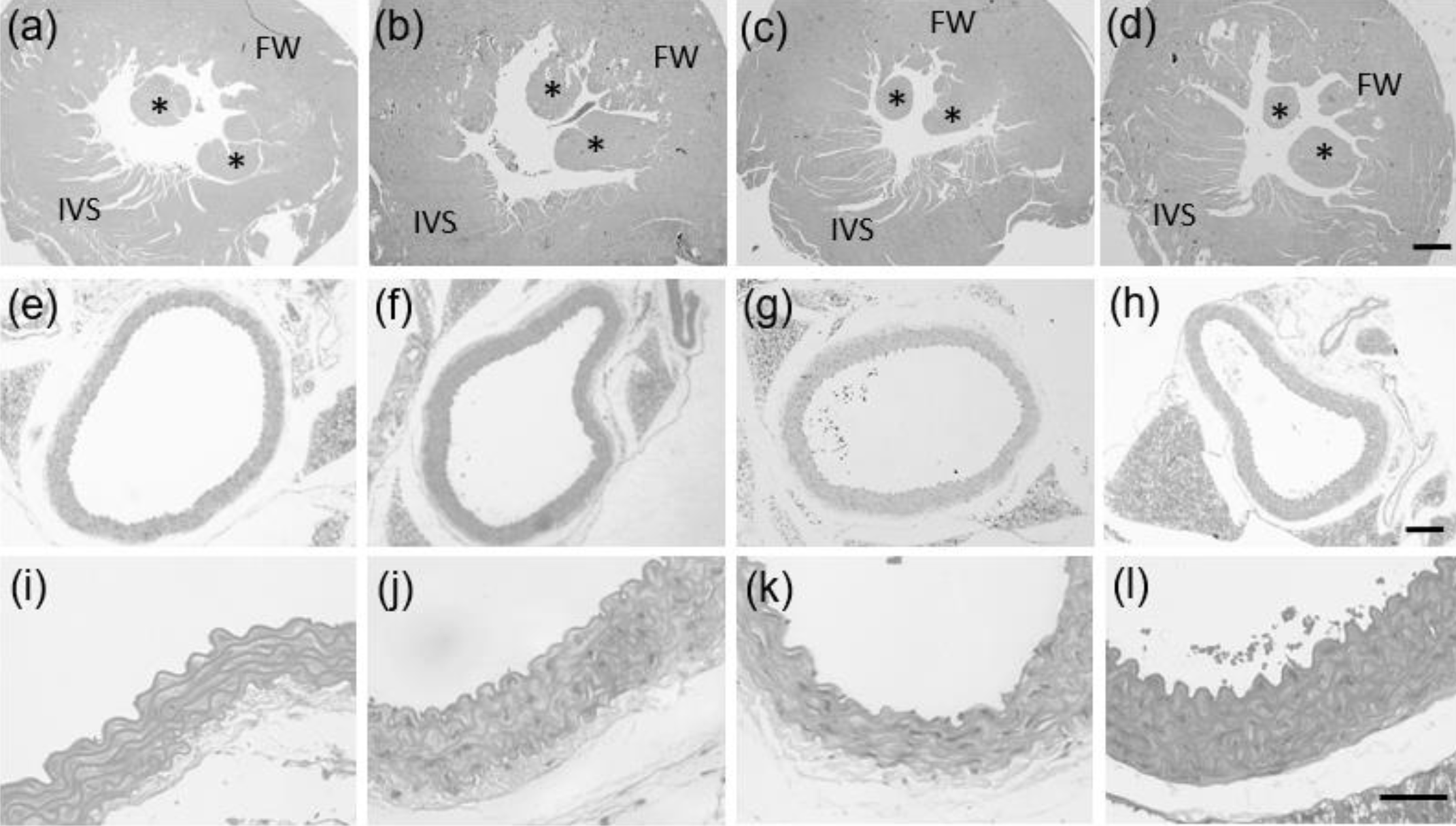
Photomicrographs of the left ventricle (**a-d**), and thoracic aorta (**e-l**). Each column represents one group (**a, e, i**: male-control; **b, f, j**: male-HFSS; **c, g, k**: female-control; **d, h, l**: female-HFSS). Calibration bars represents 500µm (**d**), 300µm (**h**), and 50µm (**l**). FW, free wall (left ventricle); IVS, interventricular septum; * papillary muscles.

Intramyocardial arteries displayed wall thickening by HFSS feeding in male mice (+92%, P<0.01, table 3) and just a trend in the female. Similar to LV, intramyocardial arteries were dilated in both male (+83%, P<0.05) and female mice (+113%, P<0.01, table 3). These data were influenced solely by diet (two-way ANOVA, table 3). The thoracic aorta also presented wall thickening, but not lumen dilation. Intima-media thickness increased 27% and 21% (P<0.0001) in male and female mice, respectively, when fed for 12 weeks with the HFSS diet (table 3 and fig. 4i-l). The ratio between wall thickness and lumen diameter also increased in both gender, being 16% (P<0.05) and 26% (P<0.01) higher in male and female mice, respectively (table 3). Two-way analysis showed that whereas diet modulates aorta wall thickness, both diet and gender influenced IMT: lumen ratio.

## Discussion

Although the reasons are not fully elucidated, it is widely accepted that the development and the impact of several diseases are gender-dependent (World Health Organization, 2015; Onat et al., 2016). Despite this, the majority of animal studies were developed in male rats or mice. Thus female physiology and its response to injury are neglected. In the face of it, it is necessary to conduct studies that focus on how gender influences the several aspects of body function in the female, such as cardiovascular physiology and metabolism.

In the present study, male and female C57BL/6J mice were fed for 12 weeks with a modified AIN93M diet rich in saturated fat, sugar, and salt to investigate the impact of gender and diet intake on cardiovascular remodeling. Surprisingly, despite increased energy consumption, mice did not gain body mass or presented metabolic abnormalities, although female mice had increased body adiposity. Rodent diets rich in fat and sugar are routinely used to induce obesity in the C57BL/6J mice, but the time required to change body mass is still a scene of much controversy, mainly because of the diet chosen. Some authors observe an early effect on body mass after four weeks on high fat diet (Lin *et al*., 2000; Fukuchi *et al*., 2004; Anunciado-Koza *et al*., 2015), whereas there are reports that longer periods (about 20 weeks) are necessary for a high-sucrose or a high-fat diet to induced obesity (Fukuchi *et al*., 2004; Hariri *et al*., 2010; Krishna *et al*., 2016).

The diet chosen to carry out the present study was based on the Surwit formulae published in 1995 to induce obesity and diabetes (Surwit *et al*., 1995). The Surwit diet was previously used by our group to induce obesity and diabetes in male C57BL/6J mice. After one week, mice already presented body mass gain, and after six weeks glucose and insulin were elevated and mice presented insulin resistance (Fernandes-Santos, Carneiro, *et al*., 2009). In the present study, the Surwit formula was modified by the addition of 8% salt to induce cardiovascular remodeling, but unexpectedly mice did not gain weight or presented glucose disturbance.

Female C57BL/6J mice presented increased visceral and subcutaneous adiposity after twelve weeks of high-fat, high-sucrose, and high-salt feeding. According to Forshee et al., overconsumption of sucrose increases visceral fat deposition, promotes hepatic lipogenesis, and lipid peroxidation (Forshee *et al*., 2007). Also, it promotes the release of very low-density lipoproteins to blood circulation, which serves as a source of fatty acids to the adipose tissue to synthesize triacylglycerol (Forshee *et al*., 2007). The increased adiposity might explain in part the improvement of glucose tolerance in HFSS-fed female mice. Adipocyte proliferation in the white adipose tissue serves as a buffering mechanism, where excessive circulating glucose is uptaken by adipocytes that converts it into triacylglycerol for storage. Since glucose is cleared from the bloodstream, the pancreas does not need to liberate a great amount of insulin, and its serum levels also decrease (Hogan *et al*., 2011; Shao e Tian, 2015). The mechanisms behind the different response between male and female mice need to be further investigated.

Regarding cardiovascular remodeling, only male mice displayed heart hypertrophy. The left ventricle was not hypertrophied in male and female mice, but its lumen was dilated. The intramyocardial arteries and the thoracic aorta of male mice presented intima-media thickening, but in the female, it was only noticed in the thoracic aorta. Finally, lumen dilation occurred only in the intramyocardial arteries of both genders, but not in the thoracic aorta. Changes in LV dimension and the arterial remodeling were influenced by both gender and the HFSS diet.

The role of salt intake on LV hypertrophy is well established. Salt overconsumption raises blood pressure and leads to local (cardiac) activation of the renin-angiotensin-aldosterone system, in which angiotensin stimulates cardiomyocyte and smooth muscle cell hypertrophy through AT1 receptor activation (Mahmud e Feely, 2004; Paul et al., 2006; Aaron e Sanders, 2013). A study conducted in C57Bl/6 mice to evaluate the gender impact on salt sensitivity increased blood pressure after one day on a 4% NaCl diet in male and female mice, and it became significantly high in the sixth day on this diet (Ma *et al*., 2017). Not only salt, but also fat overconsumption for nine weeks induce cardiac hypertrophy in male C57BL/6J mice, both alone (high-fat) or in combination with salt (high-fat high-salt diet), although at different extents (Costa et al., 2012). Additionally, it was shown that salt added to these diets might potentialize the effect of macronutrients on blood pressure (Mahmud e Feely, 2004).

A limitation of the present study is that blood pressure was not assessed. However, it is largely proved that salt overload raises blood pressure in C57BL/6J mice (Mills et al., 1993; Costa et al., 2012). In the present study, mice had increased water intake, and cardiovascular remodeling though dilation is a sign of volume overload (Shimizu e Minamino, 2016). Also, increased water intake is a compensatory mechanism that aims to remove the excess of sodium from the body, along with water, through renal glomeruli (Costa *et al*., 2012).

### Conclusion

Male and female C57BL/6J mice suffered cardiovascular remodeling after 12 weeks of high-fat, high-sucrose, high-salt feeding, although they did not develop obesity or diabetes. Sexual dimorphism occurred in response to diet for body adiposity, heart hypertrophy, and intramyocardial artery remodeling.

## Authors contribution

C.F-S conceived the work. D.C.P-S, R.P.M.S., C.C-P, and C.F-S performed the experiments and data collection. D.C.P-S, R.P.M.S., and C.F-S performed data analysis and interpretation. D.C.P-S. drafted the article, and C.F-S made a critical revision of the article. D.C.P-S, R.P.M.S., C. C-P and C.F-S approved the definitive version to be published.

## Acknowledgements

Authors are thankful for Dilliane da Paixao Rodrigues Almeida for her technical assistance.

## Conflict of interest

There are no conflicts of interest to declare.

## Funding statement

This study was supported by a grant from the Brazilian agency FAPERJ (E-26/210.525/2014).

